# dsRID: Editing-free in silico identification of dsRNA region using long-read RNA-seq data

**DOI:** 10.1101/2023.06.02.543466

**Authors:** Ryo Yamamoto, Zhiheng Liu, Mudra Choudhury, Xinshu Xiao

## Abstract

Double-stranded RNAs (dsRNAs) are potent triggers of innate immune responses upon recognition by cytosolic dsRNA sensor proteins. Identification of endogenous dsRNAs helps to better understand the dsRNAome and its relevance to innate immunity related to human diseases. Here, we report dsRID (double-stranded RNA identifier), a machine learning-based method to predict dsRNA regions *in silico*, leveraging the power of long-read RNA-sequencing (RNA-seq) and molecular traits of dsRNAs. Using models trained with PacBio long-read RNA-seq data derived from Alzheimer’s disease (AD) brain, we show that our approach is highly accurate in predicting dsRNA regions in multiple datasets. Applied to an AD cohort sequenced by the ENCODE consortium, we characterize the global dsRNA profile with potentially distinct expression patterns between AD and controls. Together, we show that dsRID provides an effective approach to capture global dsRNA profiles using long-read RNA-seq data.

## Introduction

Cytosolic double-stranded RNAs (dsRNAs), upon recognition by dsRNA sensor proteins, can trigger innate immune responses^1^. This mechanism constitutes a primary means in human cells to defend against viral infections. However, dsRNAs are also generated endogenously, many of which may be candidate binding targets of cytosolic sensor proteins, such as MDA5, RIG-I or PKR. Unwanted activation of antiviral signaling by endogenous dsRNAs is prevented at least partly by the Adenosine-to-Inosine (A-to-I) RNA editing. A-to-I editing is performed by the adenosine deaminase acting on RNA (ADAR) enzymes that bind to dsRNAs^2,3^. Accumulating evidence suggests that A-to-I editing by ADAR and its binding to endogenous dsRNA affect dsRNA immunogenicity, implicated in cancer, autoimmune and inflammatory diseases^4–6^.

Identification of endogenous dsRNAs related to immunogenicity remains a major challenge. Since ADAR is a dsRNA-binding protein, A-to-I editing sites have been used as indicators of the existence of dsRNA regions. To this end, methods have been developed to leverage editing-enriched regions (EERs) to define endogenous dsRNA structures^7,8^. This type of method may use all known editing sites, such as those cataloged in RNA editing databases^9–11^, to enable a comprehensive identification of possible dsRNAs. However, the resulting dsRNAs may not be specific to the samples under study. Alternatively, RNA editing sites identified in the samples at hand may be used in the analysis, with the risk of limited coverage as it is likely that only a subset of true editing sites are identified. Despite these limitations, EER-based methods are the gold-standard approaches in identifying dsRNAs with potential relevance to innate immunity.

dsRNA structures that undergo no or low-level RNA editing in a specific sample may escape from identification by EER-based methods^12^. Low RNA editing levels may result from regulation of ADAR activities or competition between RNA-binding proteins and ADAR^13,14^. Such unedited dsRNAs may be potent activators of antiviral signaling. Thus, it is important to design methods for editing-independent identification of dsRNAs. Experimental methods (such as SHAPE or PARS) are available for global RNA structure analysis independent of RNA editing^15,16^. In addition, protein-RNA binding profiling for dsRNA-binding proteins provides a basis to infer dsRNA regions^13,14^. However, most of the above experimental methods possess limited sensitivity due to technical challenges. Methods to detect dsRNA computationally in a high-throughput manner are highly desirable.

In this work, we developed a new approach, named double-stranded RNA Identifier (dsRID), to detect dsRNA regions in an editing-agnostic manner. This method is built upon a previous observation made by us and others that dsRNA structures may induce region-skipping in RNA-sequencing (RNA-seq) reads, an artifact likely reflecting intramolecular template switching in reverse transcription^17–20^. Leveraging this observation and long-read RNA-seq data, we constructed a machine-learning model that extracts features from mapped reads and outputs predictions of dsRNA regions. Using features related to region-skipping, dsRID achieved in-silico identification of dsRNA regions independent of editing with high accuracy. We applied this method to a few long-read RNA-seq data derived from Alzheimer’s disease (AD) and control samples, which predicted novel dsRNAs with low RNA editing levels.

## Results

### Overview of the dsRID method

In a previous study with long-read RNA-seq data, we observed that many reads contained internal skipped regions that mimic spliced-out introns^17^. However, such region-skipping is unlikely a result of splicing as they were not flanked by typical splice site sequences and the starts and ends of the skipped region did not align consistently across multiple reads (**Fig. 1A**). We hypothesized that this observation reflects reverse transcriptase (RT)-generated deletion artifacts in cDNAs. As previously reported, such artifacts may be caused by intramolecular template switching, a process where RT skips the hairpin structure of the template RNA^18–20^ (**Fig. 1B**).

**Figure 1:**
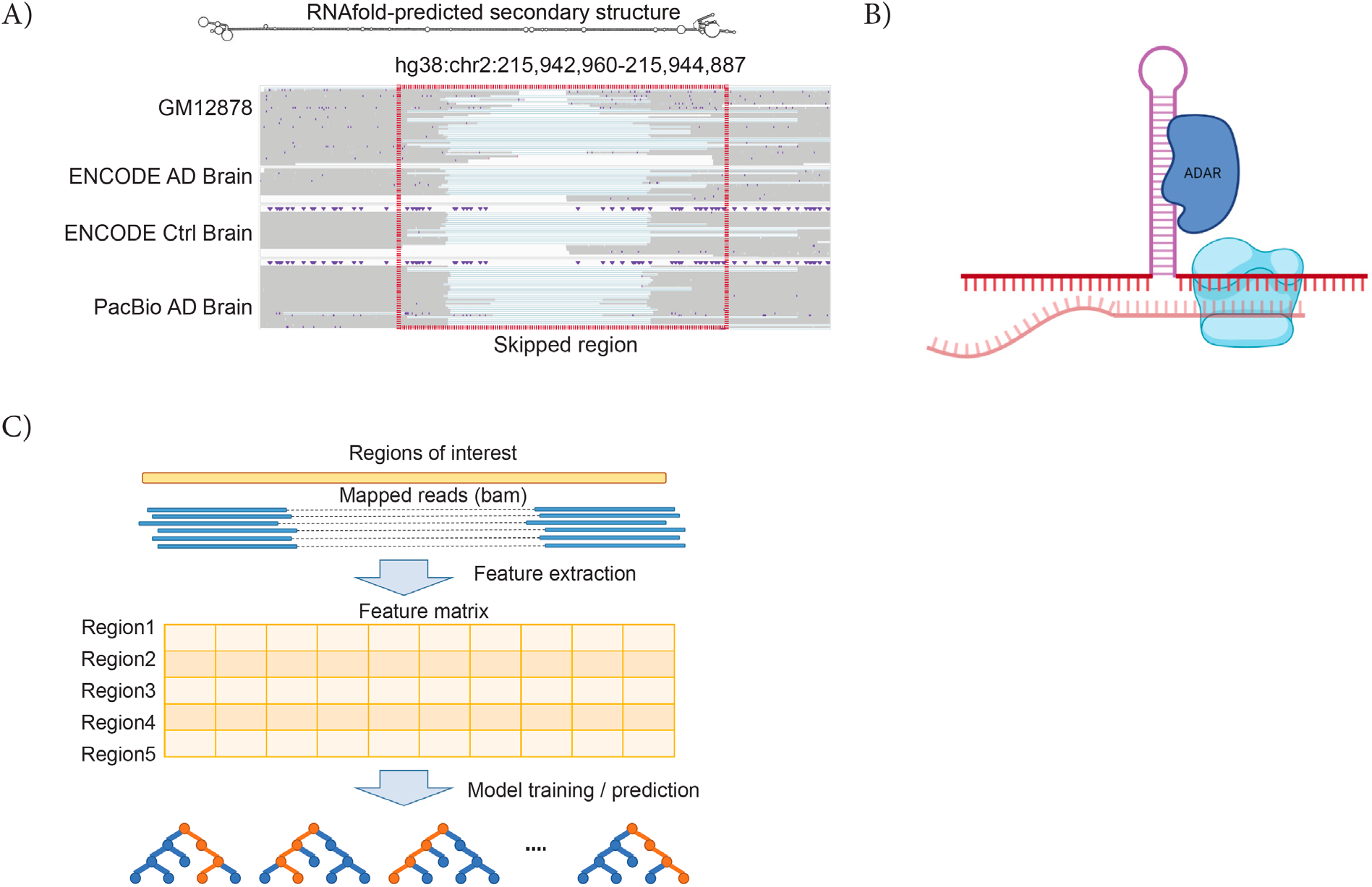
Overview of the dsRID method. A) An example region showing internal skipping that occurs in multiple datasets. Top: RNAfold-predicted structure of the genomic region. Bottom: IGV plots of mapped reads from 4 datasets. Ctrl: control B) Schematic diagram showing the hypothesis of template skipping due to double-stranded structure and ADAR binding (created by Biorender). C) Schematic diagram for the steps in dsRID.

Inspired by the above observation, we built a machine learning model, dsRID, to predict whether certain transcripts form dsRNA structures using only features related to internal region-skipping in the long reads. The dsRID method consists of three main steps: feature extraction, training and prediction. After a standard read mapping procedure using minimap2^21^, we focused on regions (2500nt in length) with at least 6 mapped reads and at least 1 read with internal skipping. We extracted a number of features from such regions, for example, skipping ratio (calculated as the ratio of reads that contained internal skipping among all reads overlapping a region), skipping length (calculated as the average length of internal skipping harbored in all reads of a region) and standard deviations of the start and end positions of the skipped region (Methods). For training purposes, we used previously curated dsRNA regions as a positive set, which were defined based on EERs (Methods)^8^, and randomly sampled regions outside of the curated dsRNA as a negative set. Note that the random controls (2500nt in length) were also required to have ≥6 mapped reads and ≥1 read with internal skipping. Thus, such controls may encompass regions with pre-mRNA splicing events.

Following feature extraction for both positive and negative sets, we trained binary classifier models such as random forests, logistic regression and support vector machines to predict dsRNA regions (**Fig. 1C**). We used TPOT^22^ to tune the hyperparameters and select the model with the best performance (Methods). In the prediction step, we applied the model to regions across the genome, excluding positive regions with curated dsRNAs (Methods). Given the binary classification problem, we defined predicted dsRNA regions as those that passed the threshold of 0.5 in the predicted probability. It should be noted that, in applying dsRID, the users can choose to skip the training step and carry out the predictions with our pretrained model. For the predicted “candidate dsRNAs”, we further applied RNAfold to evaluate their structures. We denote those with long dsRNA structures as “novel dsRNAs” (Methods) and the rest as “structured RNAs”.

### dsRID predicts dsRNA regions with high performance across several datasets

We first evaluated the performance of the model using long-read RNA-seq data derived from the brain sample of an AD patient generated by Pacific Biosciences (Pacbio AD). We trained the model on 13,469 positive regions and the same number of randomly selected negative regions. We carried out 20-fold cross-validation and observed an accuracy of 89% (**Fig. 2A**). Next, we evaluated the performance of the method applied to other long-read RNA-seq datasets. Specifically, we used 10 ENCODE datasets generated from the GM12878 cells or frontal cortexes of healthy individuals or patients with AD. The number of regions used in the training step for each dataset is shown in **Fig. S1** which approximately correlated with the sequencing depth due to the coverage requirements in defining the candidate regions. It should be noted that the performance evaluation below included all predicted candidate dsRNAs (i.e., both novel dsRNAs and structured RNAs).

**Figure 2:**
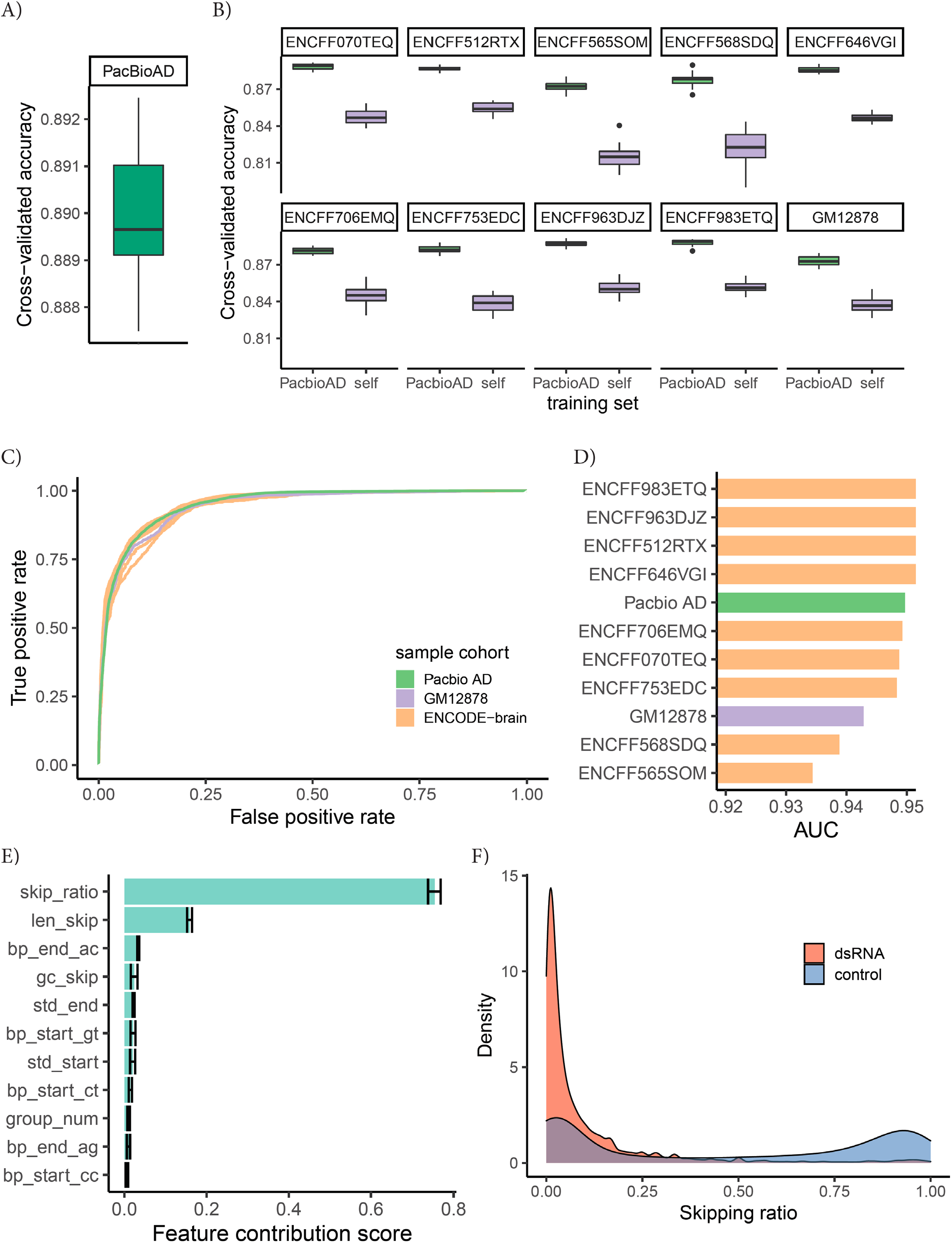
dsRID predicts dsRNA regions with high performance across several datasets. A) Box plot showing 20-fold cross-validated accuracy of dsRID trained on Pacbio-AD data. B) Box plots showing cross-validated accuracy of dsRID for different datasets. X-axis indicates whether the model is trained on its own dataset (self) or the Pacbio-AD data. C) Receiver operator curve (ROC) showing the performance of dsRID trained on the Pacbio-AD dataset. Y-axis represents true positive rate and x-axis represents false positive rate. D) Area-under-the-curve (AUC) of the ROC for each dataset. The datasets are color-coded as shown in C). E) Bar plot showing feature contribution score for each feature in the Pacbio-AD-trained dsRID model. F) Distribution of skipping ratios stratified by known dsRNA and controls.

We carried out 20-fold cross-validation for each dataset using two different models, the model trained using the same dataset and the model derived from the Pacbio AD data. When trained with each respective dataset, the average cross-validation accuracy was 84.1%. In contrast, this accuracy was 88.3% using the Pacbio AD-trained model for each dataset (**Fig. 2B**). The enhanced accuracy in the latter case likely reflects the fact that the Pacbio AD data had the highest sequencing depth and the most training regions to encompass a comprehensive dsRNA landscape (**Fig. S1**). Henceforth, we used the model trained on the Pacbio AD data for further analysis since it is the best performing model overall.

We further evaluated the performance of our model on each dataset using receiver-operator curve (ROC) analysis and calculated the area under the curve (AUC). The average AUC across all datasets was 0.95 (**Fig. 2C, D**). In addition, we used precision-recall curves to evaluate the performance and calculated the area under the precision-recall curve (AUPRC). The average AUPRC across all cohorts was 0.94 (**Fig. S2**). These results suggest that our model performs well in terms of both sensitivity and specificity.

To examine the relative importance of each feature, we performed a permutation-based feature contribution analysis (Methods). We first computed the variance explained by the model (R^2^ value). This variance was then compared to that calculated by permuting each feature vector respectively. The reduction in R^2^ upon the permutation was defined as the contribution score of the corresponding feature. We observed that the skipping ratio had the highest contribution score (75.4%), followed by the length of skipping (15.9%) (**Fig. 2E, 2F, Fig. S3**). Specifically, the positive dsRNA regions had a much lower skipping ratio than randomly sampled regions (**Fig. 2F**). This observation suggests that region-skipping due to dsRNA structures occurs randomly to a minor fraction of the cDNA molecules. Overall, the above data support the high performance of the dsRID model across multiple datasets. In addition, the model trained on the deeply sequenced data (Pacbio AD) is preferred at least for datasets tested in this study.

### Characterizations of novel dsRNA regions predicted by dsRID

A total of 82,266 candidate dsRNA regions (not present in the positive set for training) were identified across all 11 datasets (PacBio AD and 10 ENCODE datasets). As shown in **Fig. 3A**, the majority of candidate dsRNAs were unique to one dataset, which may reflect the fact that only a subset of true dsRNAs was captured in each dataset limited by sequencing depth. Alternatively (or additionally), this observation may be due to region-skipping occurring relatively randomly to structured regions. Among all candidate dsRNA regions, 32,391 were categorized as novel dsRNA based on RNAfold, and 49,875 were denoted as structured RNAs (Methods).

**Figure 3:**
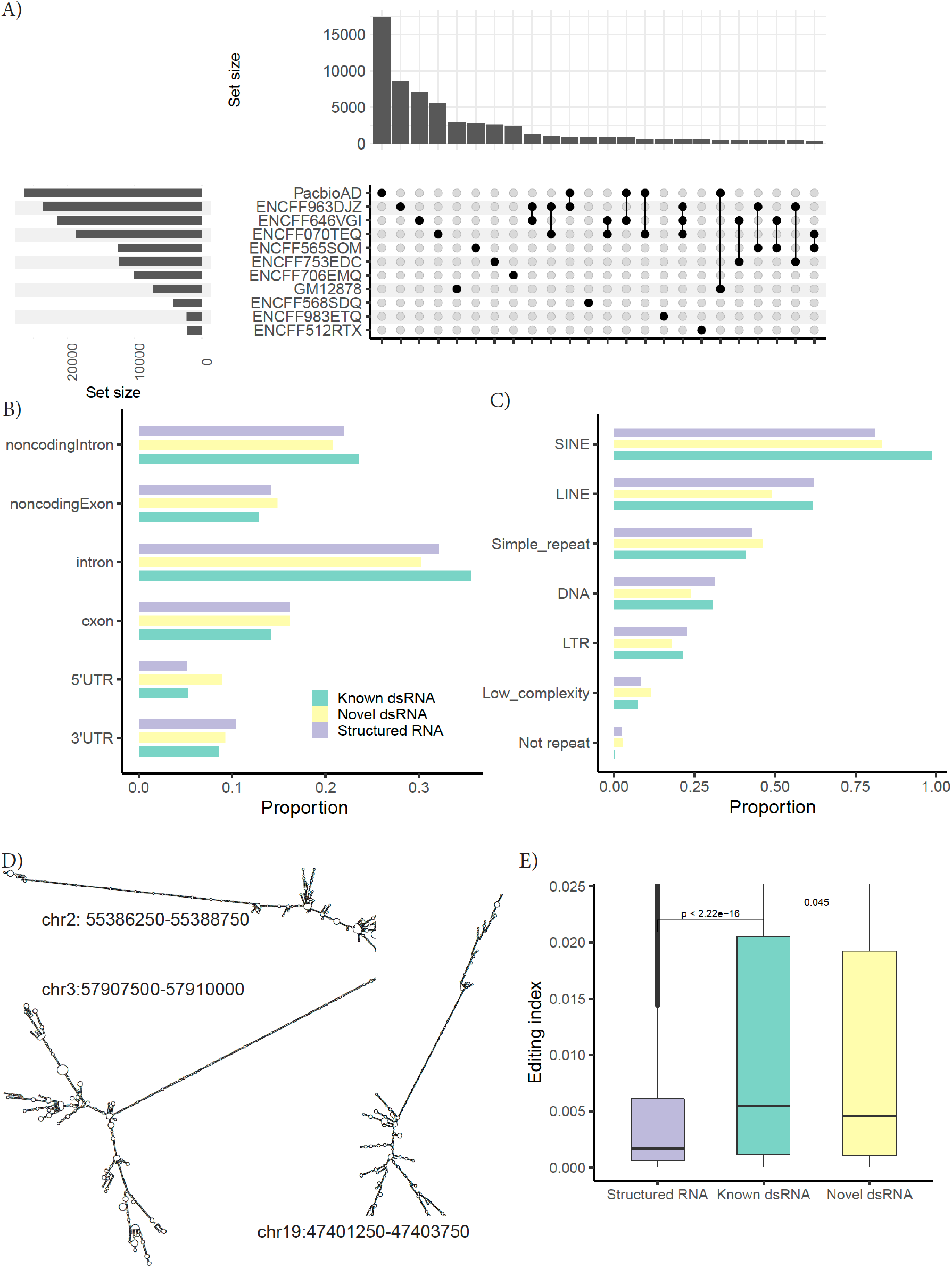
Characterization of novel dsRNA regions predicted by dsRID. A) Upset plot showing the number of novel dsRNAs detected by dsRID in each dataset and the overlaps across different datasets. Bars on the left: the number of novel dsRNAs for each dataset, bars on the top: the number of novel dsRNAs that are unique to each dataset or shared between multiple datasets. B) Proportion of known or novel dsRNAs, or structured RNAs in different types of regions. C) Proportion of known or novel dsRNAs, or structured RNAs in different types of repeats. D) Example novel dsRNA structures and their genomic coordinates (hg38). E) Editing index of known or novel dsRNA regions, or that of structured RNAs. P-values were calculated via Wilcoxon rank-sum tests.

We next analyzed the characteristics of the union of all novel dsRNAs. Similarly as known dsRNAs, novel dsRNAs most frequently overlapped with intronic regions compared to other regions. Interestingly, the novel dsRNAs were more enriched in 5’ UTRs relative to known dsRNAs (**Fig. 3B**, Proportion test, p < 2.2e-16). This observation indicates that there may exist more dsRNA structures in 5’ UTRs than appreciated previously, which may have regulatory impacts, such as translational regulation^23^. Structured RNAs did not show substantial difference in their regional distributions relative to the known or novel dsRNAs.

Furthermore, we analyzed the overlap of dsRNAs with different types of repetitive regions. As expected, most known dsRNAs overlapped with SINE elements (**Fig. 3C**), reflecting the fact that they were derived from EERs enriched in Alu regions. Although dsRID does not impose bias on the types of regions from which to discover dsRNAs, the novel dsRNAs also had high enrichment in repetitive sequences, especially SINEs, consistent with the propensity of repetitive elements forming highly structured regions. Nonetheless, compared to know dsRNAs, novel dsRNAs were significantly less enriched in SINEs (Proportion test, p < 2.2e-16), likely due to the editing-independent identification enabled by dsRID. Notably, structured RNAs also had enrichment in repetitive sequences, supporting that such RNAs had repeat-generated structures. The structures of a few example novel dsRNAs are shown in **Fig. 3D**, which harbor extended double-stranded regions. Overall, the enrichment of novel dsRNAs in repetitive sequences supports the validity of their predicted existence.

Lastly, we investigated whether the novel dsRNAs were enriched with RNA editing sites. We observed a significant but modest positive correlation between the dsRID-predicted probabilities of dsRNAs and RNA editing index (**Fig. S4**). This observation suggests that regions that are edited *in vivo* are more likely predicted as dsRNAs by our method. Nonetheless, compared to that of known dsRNAs, the RNA editing index of novel dsRNAs is slightly lower (**Fig. 3E**). However, both novel and known dsRNAs had significantly higher editing indexes than structured RNAs (**Fig. 3E**), Together, the above data support the validity of the predicted novel dsRNAs. Importantly, many novel dsRNAs discovered in this study may have low RNA editing levels, which may have been missed by previous methods built upon EERs.

### Comparative analysis of dsRNA in AD and controls detected by dsRID

To gain insights into the dsRNA profiles in AD, we conducted comparative analysis between AD and control brain samples from the ENCODE consortium. First, we asked whether the overall dsRNA (including both known and novel) profiles were distinct in AD and controls. Among all candidate regions that were tested in both AD and control samples, 76.3% were identified as dsRNAs in both groups, whereas 14.2% were specific to AD samples and 9.3% specific to controls. Proportion of AD-specific dsRNAs were significantly higher compared to control-specific dsRNAs (Fisher’s exact test, p < 2.2e-16, **Fig. 4A**). In addition, for each sample, we calculated the fraction of predicted dsRNAs and structured RNAs among all tested candidates. The AD samples showed a significantly higher dsRNA and structured RNA fractions (Wilcoxon rank-sum test, p = 0.017, **Fig. 4B**). Furthermore, the overall expression level of dsRNAs is higher in AD than in controls, suggesting higher production of dsRNAs in AD (**Fig. S5**). In contrast, the AD-specific dsRNAs had lower editing index than control-specific dsRNAs (**Fig. 4C**). The above data suggest that although the overall editing level is lower in AD samples, the total production of dsRNAs is higher in AD (**Fig. 4B, 4C, S5**).

**Figure 4:**
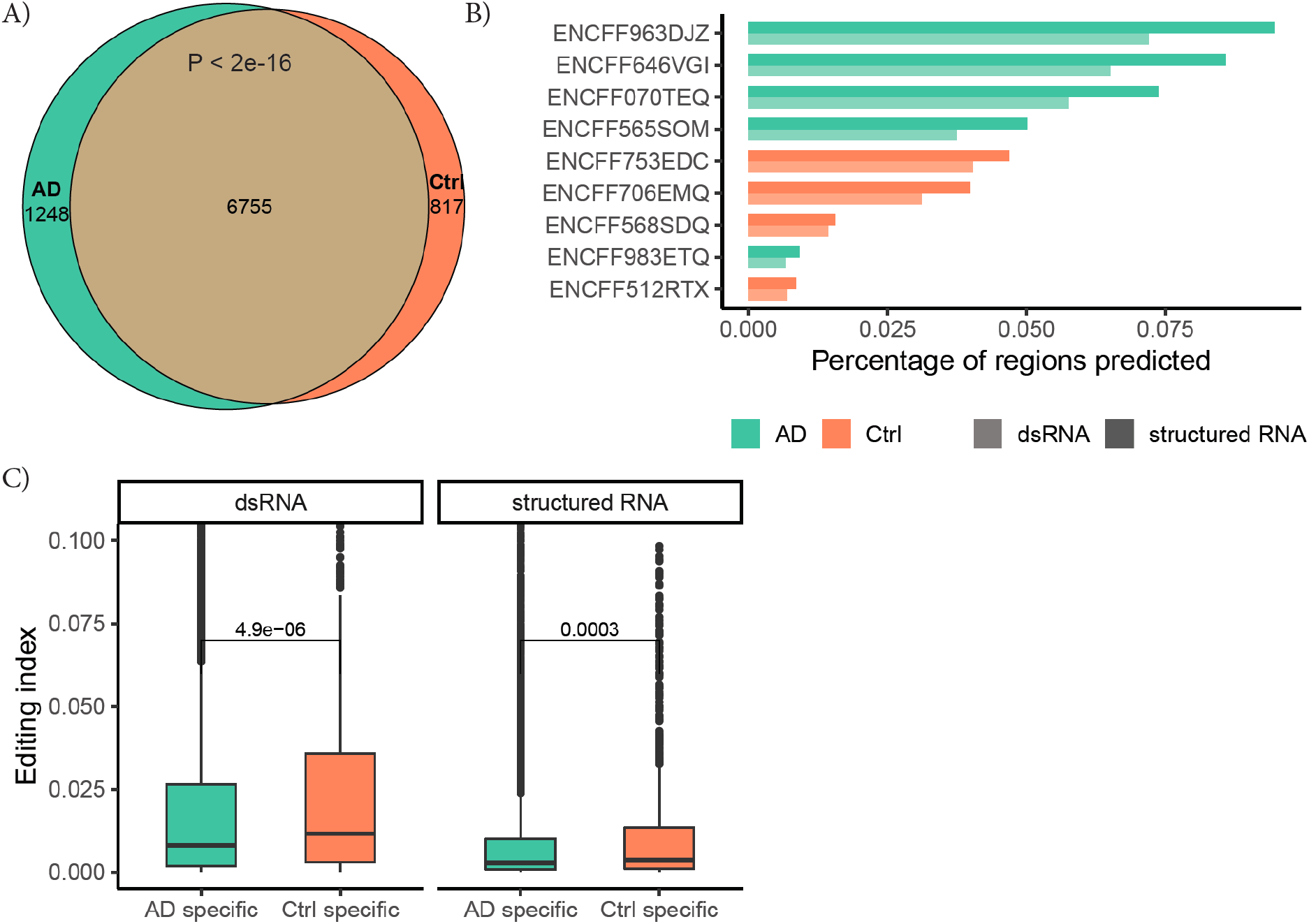
Comparative analysis of dsRNAs in AD and controls detected by dsRID. A) Venn diagram showing the overlap between dsRNAs detected in AD and control samples. B) Percentage of predicted dsRNAs or structured RNAs among all candidate regions analyzed for each dataset. C) Editing index in AD-specific and control-specific dsRNAs or structured RNA regions. P-values were calculated by two-sided t-test to compare the editing index between AD and control-specific regions.

## Discussion

Obtaining dsRNA profiles *in sillico* may greatly facilitate investigations of dsRNA-related innate immunity. In this study, we developed dsRID, a method to predict dsRNA regions using information captured in a single long-read RNA-seq dataset. The performance of dsRID is consistently high across several datasets, suggesting that the features included in dsRID reflect general characteristics of long-read RNA-seq data. dsRID identifies dsRNAs independent of RNA editing sites, in contrast to previous methods based on EERs^7,8^. We applied dsRID to data generated from AD and control brain samples. Despite the limited sample size, dsRID enabled identification of many dsRNAs, with potentially distinct expression and editing profiles between AD and controls.

Given its editing-agnostic nature, dsRID has a unique advantage over editing-based approaches in enabling dsRNA discoveries for samples with low baseline editing. Certain disease conditions, such as psoriasis^24^, autism spectrum disorders^25^, and schizophrenia^26^ are known to have reduced RNA editing levels overall. In such scenarios, identification of dsRNAs based on editing enrichment may yield limited sensitivity. dsRID makes predictions based solely on features in mapped reads, making it possible to examine the potential existence of dsRNAs outside of EERs.

Among the features used in dsRID, skipping ratio and length of the skipped region contributed the most to the model. The skipping ratios of dsRNA regions were generally lower than that of the random control regions. This observation indicates that RT-induced template switching occurs at a low frequency. It should be noted that during RNA isolation and RT, most RNA structures may have denatured and only very strong ones may remain. Thus, the dsRID method is suitable for searches of highly structured regions, such as those formed by EERs. In addition, the strongly structured regions may fold into other types of RNA structures, which may also cause template switching in RT. Nonetheless, the training step of dsRID focuses on dsRNAs formed by EERs, thus enriching for this type of RNA structures. Additionally, dsRID uses RNAfold to check for predicted structures, to further enrich for strong dsRNA structures.

Notably, the skipping ratios of the random controls (defined as random regions with at least 6 reads and at least 1 read with region-skipping) showed a bimodal distribution, similar to the distribution of exon inclusion levels of splicing. In contrast, the skipping length of dsRNAs is larger than that of random controls. For random controls, a region of 2500nt in length was considered, which is shorter than typical introns in human genes, the set of random controls may be enriched with both alternatively spliced events with relatively short introns and other skipping events due to sequencing errors/genetic variants or other reasons.

In this study, we focused on developing and applying dsRID using PacBio long-read sequencing data. In general, dsRID can be applied to data generated by other long-read sequencing technologies or short-read RNA-seq data, since the features used in the model can be derived from the other data types as well. However, the impact of different sequencing protocols on the features and performance of the method should be investigated thoroughly.

Together, we showed that dsRID is an effective method to detect dsRNA regions *in silico*. Our method featured novel dsRNA regions that are lowly edited and may be missed by EER-based approaches. Future studies highlighting long-read sequencing data in different contexts can be analyzed by dsRID to better understand the landscape of dsRNA, its regulation and function.

## Methods

### Datasets

AD long-read RNA-seq data was downloaded from PacBio (https://www.pacb.com/connect/datasets/). Long-read RNA-seq data of GM12878 cells and 9 samples of human mid-frontal cortex (AD or controls) were downloaded from the ENCODE project (https://www.encodeproject.org/ accession numbers: ENCSR962BVU, ENCSR462COR, ENCSR169YNI, ENCSR257YUB, ENCSR690QHM, ENCSR316ZTD, ENCSR697ASE, ENCSR094NFM, ENCSR463IDK, ENCSR205QMF). Reads in the fastq files were aligned using minimap2 according to the ENCODE standard parameters with the additional --cs flag for downstream analysis^21^.

### Dataset curation for training and validation

For each long-read RNA-seq dataset, we first defined positive regions and negative regions for model training. Positive regions were defined as those with known dsRNAs annotated by EER-based methods (see below)^7,17^. Negative regions were randomly sampled regions non-overlapping with the positive regions and with a window size of 2500nt. To prevent each region from having null feature values, both positive and negative regions were required to have at least 6 reads in total and at least 1 read with region-skipping. We matched the number of negative regions to the number of positive regions to balance the dataset.

### Identification of editing enriched regions

Based on the approach suggested by Whipple et al.^7^, we identified EERs using editing sites from REDIportal^9^. Regions were defined as editing enriched if there existed at least 3 editing sites in a 50 bp window. Then EERs that were within 1kb from each other were merged. These regions were then structurally verified using RNAfold^27^, and only dsRNAs with at least 200bp stem length with up to 20% of mismatches, as well as an adjusted MFE less than −0.35, were retained.

### Feature extraction

For each region of interest, we extracted features based on reads mapped to the region. The features were defined as follows:

*Skipping_ratio*: The number of reads that contained internal skipping divided by the total number of reads mapped to the region.

*Length_skipping*: Average length of skipped regions in base pairs among reads with internal skipping.

*Std_start, std_end*: standard deviation of the coordinates corresponding to the start and end sites of internal skipping, respectively.

*Gc_skip*: GC content of the skipped region

*Number of skipping groups:* The number of distinct skipping groups. The skipping start and end sites were grouped together when sites were within 100bp of each other. We assigned all sites to a group so that the left most site and right most site in each group were less than 100bp away from the median of the same group. If the number of groups for the start sites and end sites were different, we took the average number of start and end groups to denote the overall number of skipping groups from both ends.

*Bp start, end*: Two bases prior to and after each end of the skipped region, aiming to differentiate stochastic skipping from splicing that has specific splicing donor and acceptor sequences.

### Hyperparameter tuning using TPOT

We used TPOT^22^ for feature selection, model selection and hyperparameter tuning in the dsRID model trained on PacBio AD. TPOT is an automated machine learning optimization tool developed for data science applications in biomedical research.

### Permutation based feature contribution analysis

To investigate the relative contribution of each feature to the overall model, we employed permutation based feature contribution analysis using the random forest model trained with the PacBio AD dataset. First, we computed the variance explained by the model (R^2^) using a held-out validation dataset. Next, for each feature, we permuted its feature vector and recomputed R^2^ in the validation set. The decrease in the recomputed R^2^ relative to the original R^2^ was defined as the contribution score of the feature. We used the python package *scikitlearn* to conduct this procedure^28^.

### Calculation of minimum free energy for each candidate region

For each candidate dsRNA region, we used RNAfold^27^ in the Vienna RNA package to compute minimum free energy and its corresponding structure. RNAfold was run using default parameters and -AMFE flag to compute adjusted minimum free energy.

### Discovery of novel dsRNA regions

To discover novel dsRNA regions in each dataset, we analyzed windows spanning 2500nt with a sliding step of 1250nt across the genome. Those with at least 6 reads in total and at least 1 read with internal skipping were retained for feature extraction. We ran dsRID using the random forest classifier trained on the Pacbio AD dataset to compute the probability of forming a dsRNA in each region. For regions with more than 50% probability of being dsRNA and without overlapping with EER-based dsRNAs, we further examined their folded structure using RNAfold. For regions with a stem length of more than 200bp and up to 20% mismatches and an adjusted MFE less than −0.35, we denoted them as novel dsRNA regions and otherwise structured RNAs.

## Data availability

Pacbio long-read RNA-seq data is available via https://downloads.pacbcloud.com/public/dataset/Alzheimer2019_IsoSeq. Long-read RNA-seq data of GM12878 is available via the ENCODE project with accession number ENCSR462COR.. Alzheimer’s disease and control brain long-read RNA-seq data are available via the ENCODE project with accession numbers ENCSR169YNI, ENCSR257YUB, ENCSR690QHM, ENCSR316ZTD, ENCSR697ASE, ENCSR094NFM, ENCSR463IDK, ENCSR205QMF.

## Code availability

Software implementation of dsRID is available in github under https://github.com/gxiaolab/dsRID.

## Acknowledgements

We would like to thank members of the Xiao laboratory for providing helpful comments on this work. This work was supported in part by grants from the National Institutes of Health (R01MH123177, R01AG078950 to X.X.). M.C. was supported by the NIH T32LM012424.

## Reference

1. Cheng, G., Zhong, J., Chung, J. & Chisari, F. V. Double-stranded DNA and double-stranded RNA induce a common antiviral signaling pathway in human cells. Proceedings of the National Academy of Sciences 104, 9035–9040 (2007).

2. Liddicoat, B. J. et al. RNA editing by ADAR1 prevents MDA5 sensing of endogenous dsRNA as nonself. Science 349, 1115–1120 (2015).

3. Nakahama, T. & Kawahara, Y. The RNA-editing enzyme ADAR1: a regulatory hub that tunes multiple dsRNA-sensing pathways. International Immunology dxac056 (2022) doi:10.1093/intimm/dxac056.

4. Wang, H., Chen, S., Wei, J., Song, G. & Zhao, Y. A-to-I RNA Editing in Cancer: From Evaluating the Editing Level to Exploring the Editing Effects. Frontiers in Oncology 10, (2021).

5. Li, Q. et al. RNA editing underlies genetic risk of common inflammatory diseases. Nature 608, 569–577 (2022).

6. Chan, T. W., Dodson, J. P., Arbet, J., Boutros, P. C. & Xiao, X. Single-Cell Analysis in Lung Adenocarcinoma Implicates RNA Editing in Cancer Innate Immunity and Patient Prognosis. Cancer Research 83, 374–385 (2023).

7. Whipple, J. M. et al. Genome-wide profiling of the C. elegans dsRNAome. RNA 21, 786–800 (2015).

8. Blango, M. G. & Bass, B. L. Identification of the long, edited dsRNAome of LPS-stimulated immune cells. Genome Res 26, 852–862 (2016).

9. Mansi, L. et al. REDIportal: millions of novel A-to-I RNA editing events from thousands of RNAseq experiments. Nucleic Acids Research 49, D1012–D1019 (2021).

10. Ramaswami, G. & Li, J. B. RADAR: a rigorously annotated database of A-to-I RNA editing. Nucleic Acids Research 42, D109–D113 (2014).

11. Kiran, A. & Baranov, P. V. DARNED: a DAtabase of RNa EDiting in humans. Bioinformatics 26, 1772–1776 (2010).

12. Reich, D. P. & Bass, B. L. Mapping the dsRNA World. Cold Spring Harb Perspect Biol 11, a035352 (2019).

13. Quinones-Valdez, G. et al. Regulation of RNA editing by RNA-binding proteins in human cells. Commun Biol 2, 1–14 (2019).

14. Rybak-Wolf, A. et al. A Variety of Dicer Substrates in Human and C. elegans. Cell 159, 1153–1167 (2014).

15. Loughrey, D., Watters, K. E., Settle, A. H. & Lucks, J. B. SHAPE-Seq 2.0: systematic optimization and extension of high-throughput chemical probing of RNA secondary structure with next generation sequencing. Nucleic Acids Research 42, e165 (2014).

16. Kertesz, M. et al. Genome-wide Measurement of RNA Secondary Structure in Yeast. Nature 467, 10.1038/nature09322 (2010).

17. Liu, Z. et al. L-GIREMI uncovers RNA editing sites in long-read RNA-seq. 2022.03.23.485515 Preprint at https://doi.org/10.1101/2022.03.23.485515 (2022).

18. Cocquet, J., Chong, A., Zhang, G. & Veitia, R. A. Reverse transcriptase template switching and false alternative transcripts. Genomics 88, 127–131 (2006).

19. Houseley, J. & Tollervey, D. Apparent Non-Canonical Trans-Splicing Is Generated by Reverse Transcriptase In Vitro. PLOS ONE 5, e12271 (2010).

20. Tardaguila, M. et al. SQANTI: extensive characterization of long-read transcript sequences for quality control in full-length transcriptome identification and quantification. Genome Res. 28, 396–411 (2018).

21. Li, H. New strategies to improve minimap2 alignment accuracy. Bioinformatics 37, 4572– 4574 (2021).

22. Le, T. T., Fu, W. & Moore, J. H. Scaling tree-based automated machine learning to biomedical big data with a feature set selector. Bioinformatics 36, 250–256 (2020).

23. Leppek, K., Das, R. & Barna, M. Functional 5′ UTR mRNA structures in eukaryotic translation regulation and how to find them. Nat Rev Mol Cell Biol 19, 158–174 (2018).

24. Shallev, L. et al. Decreased A-to-I RNA editing as a source of keratinocytes’ dsRNA in psoriasis. RNA 24, 828–840 (2018).

25. Tran, S. S. et al. Widespread RNA editing dysregulation in brains from autistic individuals. Nat Neurosci 22, 25–36 (2019).

26. Choudhury, M. et al. Widespread RNA hypoediting in schizophrenia and its relevance to mitochondrial function. Science Advances 9, eade9997 (2023).

27. Lorenz, R. et al. ViennaRNA Package 2.0. Algorithms Mol Biol 6, 26 (2011).

28. Pedregosa, F. et al. Scikit-learn: Machine Learning in Python. Preprint at https://doi.org/10.48550/arXiv.1201.0490 (2018).

